# Unilateral resistance training induces greater rate coding adaptations in high-threshold motor units during maximal voluntary contractions

**DOI:** 10.64898/2026.06.26.734811

**Authors:** E. Lecce, P. Amoruso, A. Del Vecchio, A. Casolo, F. Felici, D. Farina, I. Bazzucchi

## Abstract

Resistance training lasting a few weeks increases maximal force mainly through neural adaptations that enhance the drive from the nervous system to muscle. While these adaptations have been well documented at the motor unit (MU) level during submaximal force contractions, the mechanisms underlying force increases during maximal voluntary contractions are poorly understood. This is due to a classic technical limitation in tracking MUs longitudinally during maximal force tasks. Here, we solved this technical challenge, enabling the investigation of MU adaptations during MVCs in both the trained and untrained limbs following unilateral resistance training. High-density surface electromyography was recorded from the biceps brachii of both limbs before and after a 4-week unilateral resistance-training intervention, and the same MUs were longitudinally tracked across sessions during MVCs by concatenation of three MVC trials of ∼5-s each.

Unilateral training increased maximal force in the *trained limb* (+16%) and induced strength transfer to the *untrained limb* (+8%). In both limbs, maximal contractions after training were characterized by greater EMG amplitude, faster muscle-fiber conduction velocity, and higher MU discharge rates, indicating enhanced neural drive to the motoneuron pool. These adaptations were strongly associated with improvements in maximal force (*R^2^* > 0.7 for all). Importantly, longitudinal MU tracking revealed a non-uniform adaptation across the MU pool: MUs with *higher baseline conduction velocity*, indicative of *higher recruitment threshold*, exhibited the largest *pre-post* increases in discharge rate, whereas lower-threshold units showed smaller changes. Collectively, these findings demonstrate that gains in maximal force and their transfer to the untrained limb are primarily mediated by enhanced rate coding of higher-threshold MUs during MVCs.

## Introduction

Resistance training (RT) is among the most widely practiced forms of exercise and increases maximal force through repeated exposure to mechanical overload (Lecce *et al*., 2026). During the first weeks of training, these gains are predominantly mediated by neural adaptations that enhance the drive from the nervous system to muscle (Škarabot *et al*., 2021). Although such adaptations have been documented at the motor unit (MU) level during submaximal contractions (Del Vecchio *et al*., 2019a; Lecce *et al*., 2025a), the MU mechanisms underlying increases in maximal force remain poorly understood. This gap is particularly important because maximal force production depends on neural strategies that differ from those typically inferred from submaximal tasks.

One useful model for studying early neural adaptations to RT is cross-education, in which unilateral training elicits measurable effects in the untrained contralateral limb without substantial changes in the contractile apparatus (Manca *et al*., 2018; Mirto *et al*., 2025). This phenomenon has often been ascribed to supraspinal modifications (Howatson *et al*., 2011, 2013; Hendy & Lamon, 2017), which in turn alter MU behaviour. Specifically, MUs in the untrained limb show increased net *discharge rate* (DR) and earlier recruitment (Lecce *et al*., 2025c), presumably due to a higher proportion of shared synaptic input to the motoneuron pool (Lecce *et al*., 2025a). Together, these changes have been associated with increased maximal strength in both the trained and contralateral untrained limbs.

To date, however, MU adaptations associated with RT and cross-education have been characterised almost exclusively during submaximal contractions. These results suggest that changes in MU recruitment, rather than DR modulation, are the principal neural mechanism associated with strength transfer to the untrained limb (Lecce *et al*., 2025c, 2025a). Yet this view may provide only a partial account of the neural adaptations relevant to strength gains, because *maximal voluntary contractions* (MVCs) rely predominantly on the discharge rate of already active MUs rather than recruitment of additional units (Heckman & Enoka, 2012). A direct assessment of MU behaviour during MVCs before and after RT is therefore needed, but has so far been prevented by a longstanding methodological limitation. Specifically, analysing MUs at maximal force is challenging because of signal complexity, overlap of MU action potentials, amplitude cancellation (Farina *et al*., 2008; Nawab *et al*., 2010; Del Vecchio *et al*., 2020), and the short duration of MVC trials, which limits the number of action potentials available for accurate decomposition (Holobar *et al*., 2014; Negro *et al*., 2016a).

When voluntary drive approaches maximal levels, several neural mechanisms contribute to the modulation of motoneuron output to muscle differently than at submaximal levels. Cortical descending drive scales with effort, reflecting increased corticospinal output and reduced intracortical inhibition (Hammond & Vallence, 2007), and is accompanied by progressive increases in *motor-evoked potential* amplitude during ramp contractions up to MVC (Castelli *et al*., 2025). Brainstem output is thought to supplement corticospinal commands and further increase the net descending excitatory input as effort rises (Glover & Baker, 2022). Notably, the relative contribution of descending pathways may change approaching the maximal voluntary force output, with evidence suggesting that reticulospinal inputs may contribute more strongly at maximal effort. In parallel, the increase in descending drive is associated with an increase in MU discharge rate (Škarabot *et al*., 2022), and this contribution appears to scale with voluntary effort (Škarabot *et al*., 2025). This suggests that MU discharge rate progressively increases to support the required force output, likely through a greater relative contribution of reticulospinal pathways, ultimately reaching its greatest effect during *maximal voluntary force* (MVF) production.

At the spinal level, MU recruitment and discharge depend on both the magnitude of synaptic input (Kernell, 1965; Schwindt, 1973; Gabriel *et al*., 2011) and its distribution across the motoneuron pool (Binder *et al*., 1998; Johnson *et al*., 2017). Intrinsic motoneuron amplification of synaptic input attributable to dendritic *persistent inward currents* (PICs) also increases as voluntary effort rises (Orssatto *et al*., 2021; Mesquita *et al*., 2024; Škarabot *et al*., 2025). This process is likely mediated by a higher monoaminergic input (Thorstensen *et al*., 2024), which exerts a proportionally greater effect at high than at low contraction intensities (Kavanagh & Taylor, 2022; Henderson *et al*., 2022).

As contractions approach MVC, most low-threshold MUs saturate their DR, so that further increases in force may depend increasingly on the rate coding of higher-threshold units (Fuglevand *et al*., 1993; Heckman & Enoka, 2012; Lecce *et al*., 2026). In some muscles (e.g., the biceps brachii), full MU recruitment occurs at ∼ 90% MVF (Kukulka & Clamann, 1981), so analyses restricted to submaximal contractions (≤ 70% MVF) may overlook mechanisms that become most relevant at maximal effort. This is particularly relevant for cross-education because, at MVC, force differences cannot be explained by recruitment alone and must instead reflect changes in rate coding. A key question is whether the overall increase in descending drive following cross-education has a different impact on low- and high-threshold MUs, for example, because of the predominant saturation of low-threshold units and the non-uniform distribution of intrinsic excitability. Since cross-education is often attributed to increased supraspinal excitability (Hortobágyi *et al*., 2011; Ruddy & Carson, 2013; Hendy & Lamon, 2017) and motoneuron output reflects both the magnitude of synaptic input and its amplification (Lee & Heckman, 2000; Farina *et al*., 2016; Lecce *et al*., 2025b), studying MVC provides a stronger test of whether these central adaptations alter the magnitude or the distribution of neural drive when the neuromuscular system operates at its maximal voluntary level.

Using HDsEMG and a slightly modified concatenation-based decomposition approach, we identified a relatively large population of MUs during *MVC trials* to characterize spinal motoneuron adaptations underlying changes in maximal force in both the trained and the contralateral untrained limbs after a 4-week unilateral RT intervention. Global EMG metrics (*muscle fiber conduction velocity* [MFCV] and EMG amplitude), which have been shown to reflect MU adaptations in *longitudinal* RT interventions (Casolo *et al*., 2020a; Del Vecchio *et al*., 2025), were also examined to provide a more reliable basis for comparison. Based on the above-mentioned evidence, we hypothesized that both the trained and untrained limbs would exhibit (*a*) increased MUDR, RMS, and MFCV, (*b*) greater MUDR changes in higher-threshold MUs, and (*c*) these neural modifications would correlate with increases in maximal muscle force.

## Methods

### Participants and ethical approval

The study was approved by the local ethics committee of the University of ‘Foro Italico’, Rome (approval *CAR157/2023*), and adhered to the standards outlined in the Declaration of Helsinki. All participants provided written informed consent, which detailed the experimental procedures and potential risks and emphasized their right to withdraw from the study at any time without consequences.

Sample size was determined a priori in G∗Power 3.1 (Faul *et al*., 2007). Considering the mixed-effect model (F-test, ANOVA repeated measure, within–between interaction, f = 0.40, α = 0.05, 1 − β = 0.80, two groups, two time points, r = 0.50), 16 participants (8 per group) were required. Given an expected withdrawal rate of 20%, we recruited 20 participants (10 males and 10 females). Following a familiarisation session, participants were randomly assigned to either the intervention group (INT, n = 10; females = 5) or the control group (CNT, n = 10; females = 5) using a block-randomization approach to ensure equal group sizes (Kang *et al*., 2008).

Each participant was assigned a unique alphanumeric code to protect their privacy and confidentiality. Exclusion criteria included metabolic and/or upper limb musculoskeletal disorders, acute infections, uncontrolled hypertension, use of medications that affect muscle protein metabolism, vascular tone, or neural activity, and use of oral contraceptives following previous longitudinal setups in resistance RT (Burrows & Peters, 2007; Elliott-Sale *et al*., 2020; Reif *et al*., 2021). Inclusion criteria required participants to be between 18 and 35 years old and in good health.

### Study overview

The study design was previously detailed (Lecce *et al*., 2025a). Briefly, participants attended three laboratory visits, the first to be informed and familiarised with experimental procedures and the subsequent two to perform the neuromuscular tests at baseline (T0) and after the 4-week RT intervention (T1) or the control period. RT consisted of 12 training sessions (three per week) over 4 weeks, with a washout period of 5-7 days before neuromuscular testing to minimize muscle soreness and residual training effects (Cheung *et al*., 2003)

The elbow-flexor force was evaluated during the experimental setup, and the biceps brachii was selected for its primary role in generating elbow flexion force (Dartnall et al., 2008; Yu, Zhang et al., 2022) for the concurrent recording of HDsEMG. Each participant’s dominant limb was identified using the *Edinburgh Handedness Inventory Questionnaire* (Oldfield, 1971). Although some evidence suggests that training a less habitually used limb may enhance cross-education (Farthing *et al*., 2007), other findings report greater transfer when the dominant limb is trained (Farthing, 2009) or no difference between limbs (Song et al., 2024). Considering this contrasting evidence, we trained the non-dominant limb to minimize bias introduced by daily activities.

No measurements were taken during the first visit. Data collection took place in the second visit (T0), during which assessments of MVF and HDsEMG recordings were obtained from the biceps brachii of both limbs, one limb at a time (Fig. 1). The same data collection was repeated following the intervention or control period (T1).

**Figure 1.**
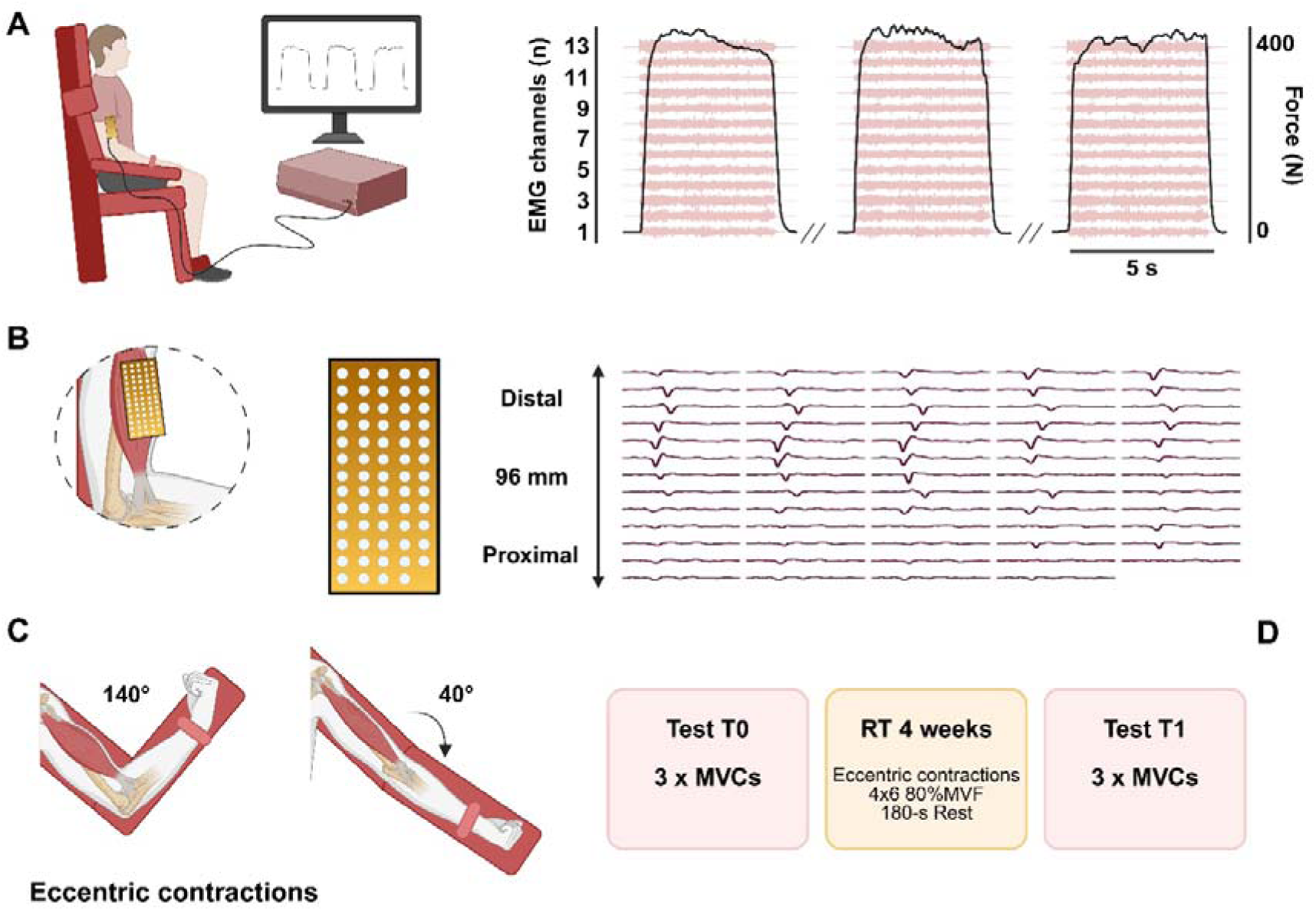
Study overview and experimental design. (A) experimental setup for neuromuscular tests composed of at least three maximal voluntary contractions (MVCs) concatenated prior to decomposition. The force trace (black line) was synchronized with the EMG signal. (B) A bi-dimensional electrode grid (13 rows × 5 columns) was placed over the biceps brachii to record myoelectrical activity during the MVC trials. (C) The training intervention comprised eccentric contractions of the elbow flexors at the isokinetic dynamometer (Kin-Com). (D) A summary of the experimental design, including timelines, is reported in boxes. The image was created with *Biorender* (https://app.biorender.com).

Since the influence of menstrual cycle-related fluctuations on MU discharge properties has been evidenced (Piasecki *et al*., 2023; Jenz *et al*., 2026), female participants not using hormonal contraception were tested during a predefined menstrual phase to minimize hormone-related variability in neural excitability and MU characteristics (Tenan *et al*., 2013; Weidauer *et al*., 2020; Piasecki *et al*., 2023). This included either the ovulatory or mid-luteal phase, identified with a validated *Menstrual Practices Questionnaire* (Hennegan *et al*., 2020). These phases were chosen because the low-threshold MU firing rates during ovulation and the mid-luteal phase are similar, allowing standardization of testing under conditions of heightened hormonal influence while avoiding the variability associated with the early follicular phase (Piasecki *et al*., 2023). Post-intervention testing was conducted during the same menstrual phase as baseline to maintain consistency.

### Experimental procedures

The volunteers were instructed to refrain from strenuous exercise and caffeine consumption for 24-48 h before testing (Amoruso *et al*., 2026). Neuromuscular testing began with a standardized warm-up consisting of seven isometric submaximal elbow flexions of approximately 3-5 s each (3 x 50%, 3 x 75%, and 1 x 90% of the perceived MVF), separated by 30 s of rest. Following the warm-up, participants performed MVCs lasting ∼ 5 s and were instructed to push *as hard as possible*, with strong verbal encouragement to ensure optimal performance (Romdhani *et al*., 2024; Lecce *et al*., 2025d). Three trials were completed, separated by 180 s of rest, and a fourth trial was conducted if the first two differed by more than 5% (Škarabot *et al*., 2024). The highest instantaneous force achieved across trials was taken as the MVF.

The holding-phase inclusion after the force rising phase was chosen based on previous studies assessing high-intensity contractions with concurrent HDsEMG recording (Del Vecchio *et al*., 2019b; Škarabot *et al*., 2022, 2024). Although these setups investigate ballistic contractions at approximately 80%MVF, we included an extended hold phase to prolong contraction duration and ensure a sufficient number of independent sources [i.e., action potentials] for reliable MU decomposition (Holobar *et al*., 2014). Trials with countermovement or pre-tension were excluded and repeated (Škarabot *et al*., 2024). During experimental sessions, participants received visual feedback of force trajectories from a monitor positioned 1.5 m from their eyes.

The intervention group performed a 4-week unilateral RT protocol at the isokinetic dynamometer (Kin-com, Chattanooga, TN, USA), with three sessions per week using the non-dominant limb. Each session began with a standardized warm-up, including three sets of 10 dynamic contractions at 30% MVF, with 60 s of rest between sets. This was followed by two sets of four eccentric contractions at 50% MVF, with 120 s of rest between sets. Subsequently, participants performed four sets of six elbow-flexor eccentric contractions at 30° s^−1^ (from 140° to 40° of flexion) at 80% MVF, with 180 s of rest between sets.

Eccentric contractions have been shown to elicit greater adaptations than other contraction regimens during the first weeks (Hortobágyi *et al*., 1996), likely attributable to distinct neural control strategies, including the preferential recruitment of high-threshold MUs that maximize force output (Enoka, 1996). Moreover, eccentric actions have been shown to induce greater cross-education effects compared with other contraction modes (Hortobágyi *et al*., 1997) and were therefore selected for the protocol. Control group participants received no intervention and were instructed to maintain their habits throughout the experimental period. This approach ensured that no behavioral modifications occurred, allowing any observed changes in the experimental groups to be attributed solely to the intervention.

### Force and HDsEMG recordings

Elbow-flexion force was measured with an isokinetic dynamometer (*load cell*, Kin-com). Participants were seated in the dynamometric chair and stabilized with chest and waist straps to minimize extraneous movement. The upper arm was positioned parallel to the trunk, while the forearm was oriented midway between supination and pronation, maintaining a 90° elbow flexion angle. The center of rotation of the lever arm was aligned with the lateral humerus epicondyle, and the wrist was secured in a cuff attached to the load cell to ensure force isolation. The analog force signal was amplified and sampled at 2048 Hz using an external analog-to-digital (A/D) converter (EMG-400, OT-Bioelettronica, Turin, Italy) to ensure synchronization with the EMG.

EMG recordings were conducted one limb at a time, randomizing among participants, using a 64-electrode adhesive grid [13 rows × 5 columns; gold-coated; electrode diameter: 1 mm; inter-electrode distance (IED): 8 mm; GR08MM1305, OT-Bioelettronica]. Following skin preparation, which included shaving, light abrasion, and cleansing with 70% ethanol, the biceps brachii long head perimeter was identified through palpation and marked with a surgical pen. The grid orientation was determined from preliminary recordings with a 16-electrode array (IED: 5 mm, OT-Bioelettronica), allowing identification of the *innervation zone* (IZ) to estimate fiber direction. The IZ was located by identifying the inversion point in the action potential propagation direction, both proximally (towards the biceps brachii proximal tendon) and distally (towards the distal tendons), along the electrode column.

The grid was positioned directly over the IZ on the muscle belly using a disposable biadhesive with layer holes adapted to the HDsEMG grids (SpesMedica, Genoa, Italy). These holes were filled with conductive paste (SpesMedica) to ensure optimal skin-electrode contact. To ensure consistent electrode positioning across all assessment time points, anatomical landmarks and skin marks were traced onto individual acetate sheets during the first assessment session (Lecce *et al*., 2025a). Reference electrodes were placed on the ulna (styloid process) and the acromion skin surface. HDsEMG signals were recorded in monopolar mode and digitized using a 16-bit multichannel amplifier (EMG-Quattrocento, OT-Bioelettronica). Signals were amplified (×150), sampled at 2048 Hz, and band-pass filtered (10-500 Hz, fourth-order Butterworth) by the EMG amplifier to remove direct current (DC – baseline offset) and aliasing artifacts while preserving the full spectral content of MU action potentials before being stored for offline analysis (Lecce *et al*., 2025a).

### Data processing

The force signal was converted to newtons (N), and gravity compensation was used to remove the offset. MVF was defined as the peak recorded force, and a 500-ms window centered on this value (+/-250 ms from MVF) was identified for EMG analysis (Fig. 2), as described below. The force signal was low-pass filtered with a fourth-order, zero-lag Butterworth filter with a cutoff frequency of 15 Hz. Only contractions without any countermovement action or pre-tension were analyzed.

**Figure 2.**
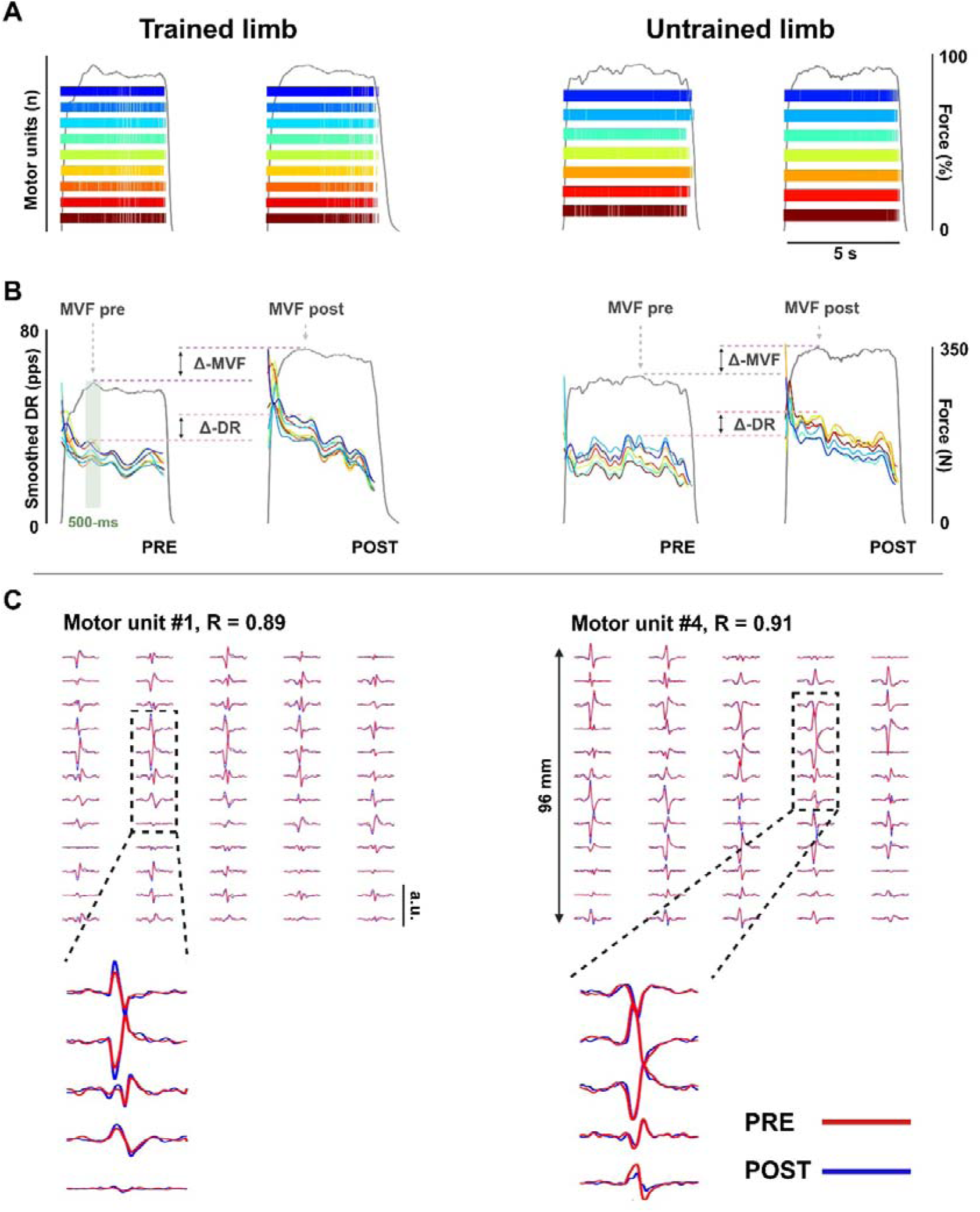
Data processing. Color-coded raster plots display MU spike trains for the identified, longitudinally tracked MUs in trained and untrained limbs (A). The smoothed discharge rate was obtained from the tracked MUs pre-intervention (T0) and post-intervention (T1), with the level of DR calculated over the *500-ms window* centred on the MVF (i.e., the best trial of the session) in both limbs (B). Tracking procedures performed to match the same MUs across the whole identified sample based on the cross-correlation of two-dimensional action potential waveforms (C). The image was created with *Biorender*.

Before processing and decomposing the raw HDsEMG signals into individual MU action potentials using *convolutive kernel compensation* (CKC) and following the amplifier hardware band-pass filtering, the monopolar EMG recordings were further band-pass filtered at 20-500 Hz with a second-order, zero-lag Butterworth filter to suppress residual movement artifacts and baseline drift (Lecce *et al*., 2025a). This two-stage approach, broad hardware filtering followed by tighter software filtering, maximizes artifact rejection and drift suppression while preserving MU action potential waveform integrity, ensuring accurate MU decomposition (Holobar *et al*., 2014).

### Global HDsEMG analysis

Double-differential signals were computed along the presumed fascicle direction (i.e., along electrode columns) to enhance the spatial selectivity of propagating MU action potentials as previously indicated (Del Vecchio *et al*., 2017, 2018c; Casolo *et al*., 2020a). The resulting signals were visually inspected, and the 4-8 channels demonstrating the clearest and most consistent propagation patterns across adjacent electrodes were selected for further analysis. EMG amplitude (µV) was quantified from the selected channels with the *root-mean-square* (RMS) extracted from the 500-ms window centered at MVF (Besomi *et al*., 2020; Del Vecchio *et al*., 2025). For each condition, the final EMG amplitude was calculated as the mean RMS across the selected channels.

Double-differential HDsEMG signals were computed from monopolar recordings along each column of the bidimensional array. Signals were visually inspected, and at least four channels per column showing clear motor unit action potential propagation from the innervation zone to the distal tendon, without waveform distortion, were selected based on a *cross-correlation coefficient* (CC) ≥ 0.85. The three central columns of each array, which consistently exhibited the highest signal quality, were retained for analysis. MFCV was then estimated within the same 500-ms window at MVF using a validated multichannel algorithm (Farina *et al*., 2001, 2004) with demonstrated high reliability (ICC ≥ 0.88) (Martinez-Valdes *et al*., 2016). The use of ≥ 4 EMG channels has been reported to detect changes in MFCV as small as 0.1 m · s^-1^ (Farina *et al*., 2002; Del Vecchio *et al*., 2018b).

### Decomposition and MU analysis

Offline decomposition and editing were performed using DEMUSE, working in MATLAB (MathWorks Inc., Natick, MA, USA). A minimum of three MVC trials (each lasting 5 seconds) were concatenated for each limb and time point before being decomposed to ensure a ∼ 15-s duration of active MU discharge activity (Fig. 1). We noticed that by concatenating at least 3 MVC trials, it was possible to accurately decode a substantially greater number of MUs from MVC than from shorter data. Moreover, by applying the separation filters obtained for the decomposed MUs at the initial MVC trials, it was possible to identify the same MUs longitudinally (i.e., across different time points/sessions) in other MVC trials (Fig 2), as long as the total interval of data after concatenation of trials was at least approximately 15 s. An experienced operator manually analyzed all identified MUs, following procedures extensively described in previous studies (Del Vecchio *et al*., 2020; Valli *et al*., 2024).

Manual editing involved inspecting and editing MU spike trains after automatic decomposition, ensuring accuracy by discarding MUs with a *pulse-to-noise ratio* (PNR) below the reference threshold (30 dB) or with discharge times separated by more than 2 s, which were excluded from analysis (Del Vecchio *et al*., 2020; Lecce *et al*., 2025b). MU duplicates, defined as units sharing at least 30% of the same discharge activity, were identified and removed using a firing-matching tolerance of 0.5 ms, with the MU having the lower PNR discarded (Škarabot *et al*., 2023).

The average MU discharge rate was computed from the series of discharge time intervals identified across the MVF 500-ms window (MUDR_MVF_) and for the entire contraction (MUDR_MEAN_). MUDR_MEAN_ and MUDR_MVF_ were averaged per participant to obtain the individual DR_MEAN_ and DR_MVF_ for linear regression analyses (see *statistical analysis*).

To reliably characterize adaptations to the RT protocol, MUs were longitudinally tracked across the intervention (T0-T1). Reported changes, therefore, reflect within-unit differences in discharge characteristics of the same MUs, avoiding bias that can occur when analyses include units identified at either time points (Fig. 2). This tracking method, based on the cross-correlation of two-dimensional action potential waveforms (Martinez-Valdes et al., 2017), has been widely used in interventional studies to assess the same set of MUs (Del Vecchio, Casolo et al., 2019; Lecce, Conti et al., 2025; Orssatto et al., 2023). Only MUs with a high cross-correlation coefficient (cutoff, R > 0.8) were included in the analysis. To further confirm the reliability of the concatenation-based decomposition, we computed the two-dimensional cross-correlation between the decomposed MU action potentials across the MVC trials used for the concatenated decomposition (Fig. 2). Specifically, MUs identified from the concatenated signals were verified to represent the same physiological unit across all individual MVC trials. This additional within-session tracking ensured that MUs derived from the concatenation approach were accurately decomposed.

MU conduction velocity (MUCV) was also computed on tracked MUs. MU action potential waveforms were extracted by spike-triggered averaging of HDsEMG signals, using the discharge times identified by decomposition (Del Vecchio *et al*., 2018c). For each MU, the first 20 discharge timings were used, with averaging performed over 15-ms intervals corresponding to MUAP duration (Casolo *et al*., 2020a). The resulting bipolar channels were first normalized to unit amplitude to minimize the influence of between-channel amplitude variability due to differences in skin-electrode contact. Double-differential signals were then derived from normalized, averaged monopolar MUAPs along the electrode columns and used to estimate MUCV. Double-differential channels were visually inspected with a customized MATLAB script, and four to eight channels from the same electrode column were selected based on the clearest action potential propagation, minimal MU shape distortion, and the highest interchannel correlation (CC ≥ 0.70) (Del Vecchio *et al*., 2018a). Because channel number affects MUCV accuracy, the largest set of channels meeting this criterion was retained. MUCV was then calculated using a multichannel maximum-likelihood algorithm, and the same number and locations of selected channels were kept for the post-intervention assessment (Farina *et al*., 2001). This analysis was performed on the same 500-ms window used for DR_MVF_ (Fig. 2) to minimize the influence of changes in discharge rate during the initial contraction phase and to standardize the discharge frequency (Campanini *et al*., 2009).

All the above procedures were performed in accordance with recent analysis guidelines (Del Vecchio *et al*., 2020; Gallina *et al*., 2022; Martinez-Valdes *et al*., 2023; Lecce *et al*., 2025b) using custom-written MATLAB scripts. Although these procedures have been less frequently applied to MVC recordings, the reliability of the decompositions in the present study is supported by the consistently high PNR index, which reflects accurate MU identification (Holobar *et al*., 2014). The tracking procedure itself provides further support for this approach, as decompositions across different sessions were performed independently, and MUs were matched only when highly consistent MUAP spatial profiles were identified across the full electrode grid (Martinez-Valdes *et al*., 2017).

### Statistical analysis

The Shapiro-Wilk test was used to assess the normality of the data. Baseline between-group differences in anthropometrical characteristics, number of identified MUs, and baseline MVF across dominant and non-dominant limbs were assessed with independent Student’s t tests.

Reliability and *agreement* in measurements were evaluated for tracked MU properties (MUCV, MUDR_MEAN_, and MUDR_MVF_) for the CNT group using two-way mixed-effects *intraclass correlation coefficients* (ICC_3,1_) using the *SimplyAgree* R-package, jamovi module. These analyses were employed to determine whether the signal processing and the decomposition process provide reliable absolute values despite the presence of an intervention (Ten Hove *et al*., 2024).

A *generalized mixed model* was used to assess the between-within-group comparisons for MVF, RMS, and MFCV in trained and untrained limbs. Between-within group comparisons of MU adaptations refer only to longitudinally tracked units and were performed for MUDR_MEAN_, MUDR_MVF_, and MUCV for both limbs separately and were analyzed using mixed-effects linear regression, an approach preserving intra- and inter-participant variability by incorporating the entire sample of extracted MUs from the 20 participants (Boccia *et al*., 2019; Yu *et al*., 2022; Wilkinson *et al*., 2023). Fixed effects of *group* and *time* and their interaction were computed, with a random intercept for each participant [e.g., MUDR_MEAN_ ∼ group × time + (1 | Participant ID)], as indicated elsewhere (Lecce *et al*., 2025b). To compare the effect of intervention across the INT group (*trained limb* [TL] and *untrained limb* [UL]) and the CNT group (*non-dominant limb* [ND] and *dominant limb* [D]), a mixed-effect model was performed on ΔMVF, ΔRMS, ΔMFCV, ΔDR_MEAN_, and ΔDR_MVF_, with a fixed effect of *limb* and the random intercept for each participant. The significance of the fixed effect was assessed by an *F-test* using Satterthwaite’s method to approximate the degrees of freedom. A *Gamma* distribution and a *Log* link function were used for modeling non-normal, positively skewed data (Ng & Cribbie, 2017). A *Bonferroni-Holm* correction was applied when needed to control for multiple comparisons in post hoc analyses, and estimated marginal means with 95% confidence intervals were computed for pre- and post-intervention comparisons.

The strength of the association (R^2^) between the change in MVF and the corresponding changes in MUDR_MEAN_, MUDR_MVF_, RMS, and MFCV for both the trained and untrained limbs was computed. Changes in DR_MVF_ were analyzed as a function of baseline MUCV to determine whether greater increases in DR occurred in MUs with higher baseline conduction velocities. In addition, the association between changes in DR_MVF_ and relative changes in MUCV was assessed to determine whether increases in MUCV were associated with concomitant increases in DR. The strength of the association was interpreted as follows: 0-0.1, *very weak*; 0.1-0.3, *weak*; 0.3-0.5, *moderate*; 0.5-0.7, *strong*; 0.7-1.0, *very strong*. To infer whether the neuromuscular adaptations imply a transfer to the contralateral untrained side, the association between the change in RMS, MFCV, MUDR_MEAN_, and MUDR_MVF_ between limbs was also computed.

Statistical analyses were performed using jamovi 2.3.28 (The jamovi project, Sydney, Australia) and SPSS 25.0 (IBM Corp., Armonk, NY, USA). A *P*-value of < 0.05 was considered statistically significant. The full statistical report, which includes INT and CNT results, is available in the supplementary material.

## Results

### Participants’characteristics

No differences in baseline characteristics and anthropometrics were observed between the INT and CNT groups (*p* > 0.05). The details of these comparisons are presented in Table 1.

**Table 1:** participants’ characteristics at baseline.

| Variable | INT | CNT | *p* |
| --- | --- | --- | --- |
| Height (cm) | 167.8 ± 8.9 | 172.8 ± 7.5 | 0.19 |
| Weight (kg) | 61.8 ± 11.5 | 70.0 ± 19.3 | 0.26 |
| BMI (kg m<sup>-2</sup>) | 21.8 ± 2.5 | 23.1 ± 4.8 | 0.45 |
| Age (years) | 23.8 ± 2.3 | 23.4 ± 2.2 | 0.69 |
| D MVF (N) | 262 ± 114 | 291 ± 115 | 0.53 |
| ND MVF (N) | 247 ± 103 | 257 ± 95 | 0.82 |
| Abbreviations: CNT, control group; D, dominant limb; INT, intervention group; ND, non-dominant limb |  |  |  |

### Decomposition and MU distribution

From the decomposition process, a total of 655 MUs were identified across the assessed MVC trials; specifically, 333 were identified in the INT group and 322 in the CNT group, with no significant difference between groups (*p* = 0.574). In the INT group, 167 MUs were found in the TL (T0: 8.8 ± 3.2; T1: 7.9 ± 2.0 per participant) and 166 in the UL (T0: 8.1 ± 1.9; T1: 8.5 ± 1.8 per participant). Of these, 52 MUs were tracked in TL (32%; 5.2 ± 1.3 per participant) and 49 were tracked in UL (29%; 4.9 ± 0.9 per participant). In the CNT group, 155 MUs were found in the ND (T0: 8.3 ± 1.9; T1: 7.2 ± 1.5 per participant) and 167 in the D (T0: 8.8 ± 2.9; T1: 7.9 ± 2.0 per participant). 48 MUs were tracked in ND (31%; 4.8 ± 1.1 per participant) and 43 were tracked in D (26%; 4.3 ± 0.8 per participant). No differences were observed across groups and conditions (*p* > 0.05). The decomposition details are presented in Table 2.

**Table 2:** Distribution of identified and tracked Mus.

| **INT** | **T0** | **T1** | **TOTAL** | **TRACKED** |
| --- | --- | --- | --- | --- |
| TL | 88 | 79 | 167 | 52 (32%) |
| UL | 81 | 85 | 166 | 49 (29%) |
| SUM | 169 | 164 | 333 | 101 (30%) |
| **CNT** | **T0** | **T1** | **TOTAL** | **TRACKED** |
| ND | 83 | 72 | 155 | 48 (31%) |
| D | 88 | 79 | 167 | 43 (26%) |
| SUM | 171 | 151 | 322 | 91 (28%) |
| **TOTAL** | **340** | **315** | **655** | **192 (29%)** |

Abbreviations: CNT, control group; D, dominant limb; INT, intervention group; ND, non-dominant limb; TL, trained limb; UL, untrained limb

### Decomposition reliability

Reliability analysis of tracked MU properties across sessions was performed only for the CNT group. Indeed, this was the group of subjects in which we could exclude changes in MU properties. The ICC values for comparisons of longitudinally tracked MUs in the CNT group are presented in Table 3. All comparisons showed good to excellent test–retest reliability, reflecting the robustness of the employed methods and tracking procedures for MVCs. To further evaluate the specificity of the tracking approach, we also computed ICC values using the full sample (i.e., non-tracked) MUs across sessions in the CNT group. In this case, reliability was markedly lower (ICC = 0.48), indicating only moderate agreement. This finding confirms that reliability estimates are substantially reduced when motor units are not consistently identified across sessions and supports the validity of the decomposition and tracking procedures, which enable reliable and accurate longitudinal identification of the same MUs over time.

**Table 3:**
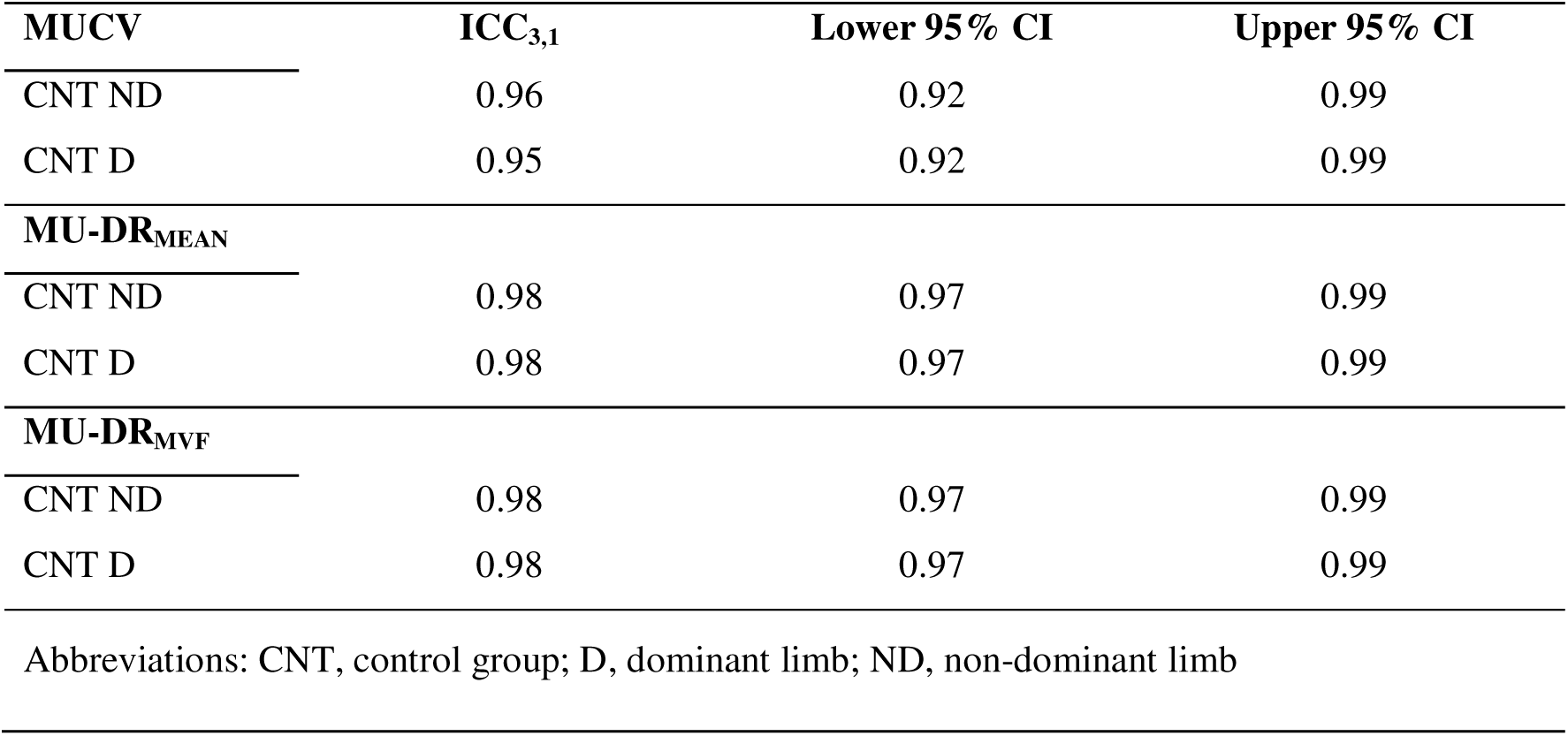
Test-retest reliability of tracked MU characteristics.

| **MUCV** | **ICC<sub>3,1</sub>** | **Lower 95% CI** | **Upper 95% CI** |
| --- | --- | --- | --- |
| CNT ND | 0.96 | 0.92 | 0.99 |
| CNT D | 0.95 | 0.92 | 0.99 |
| **MU-DR<sub>MEAN</sub>** |  |  |  |
| CNT ND | 0.98 | 0.97 | 0.99 |
| CNT D | 0.98 | 0.97 | 0.99 |
| **MU-DR<sub>MVF</sub>** |  |  |  |
| CNT ND | 0.98 | 0.97 | 0.99 |
| CNT D | 0.98 | 0.97 | 0.99 |

Abbreviations: CNT, control group; D, dominant limb; ND, non-dominant limb

### MVF

There was a significant *group* × *time* interaction for both the TL (*X²* = 67.7, *p* < 0.001) and UL (*X²* = 19.6, *p* < 0.001) for MVF. *Post-hoc* tests showed that these changes were restricted to the INT group, with a 16% higher MVF in TL (ΔMVF = 48 N, *p* < 0.001) and in UL by 8% (ΔMVF = 24 N, *p* < 0.001), while no significant differences were observed for the CNT group (*p* > 0.05). Mixed-effect model for delta results indicates a significant *limb* effect (*F_3,21.8_* = 18.2, *p* < 0.001), with *post-hoc* comparisons revealing larger gains in TL than in UL (ΔMVF = 24 N, *p* = 0.01), ND (ΔMVF = 51 N, *p* < 0.001), and D (ΔMVF = 50 N, *p* < 0.001). UL increases were also greater than in ND (ΔMVF = 27 N, *p* = 0.01) and D limbs (ΔMVF = 26 N, *p* = 0.007).

### MFCV and RMS

For RMS, a significant *group* × *time* interaction was found for TL (*X²* = 10.4, *p* = 0.002 [Fig. 3A]) and UL (*X²* = 7.6, *p* = 0.006 [Fig. 3A]). *Post-hoc* comparisons showed higher RMS confined to the INT group, in both TL (ΔRMS = 137 μV [+ 10%], *p =* 0.01 [Fig. 3B]) and UL (ΔRMS = 179 μV [+10%], *p* < 0.001 [Fig. 3B]), with no differences found in the CNT group (*p* > 0.05 [Fig. 3B]). The mixed-effect analysis indicated a significant *limb* effect (*F_3,26.1_* = 6.9, *p* = 0.001), with greater delta-RMS in TL than ND (ΔRMS = 95 μV, *p* = 0.02 [Fig. 3B]) and D limbs (ΔRMS = 122 μV, *p* = 0.04 [Fig. 3B]) and in UL compared to ND (ΔRMS = 138 μV, *p* = 0.01 [Fig. 3B]) and D limbs (ΔRMS = 164 μV, *p* = 0.002 [Fig. 3B]).

**Figure 3.**
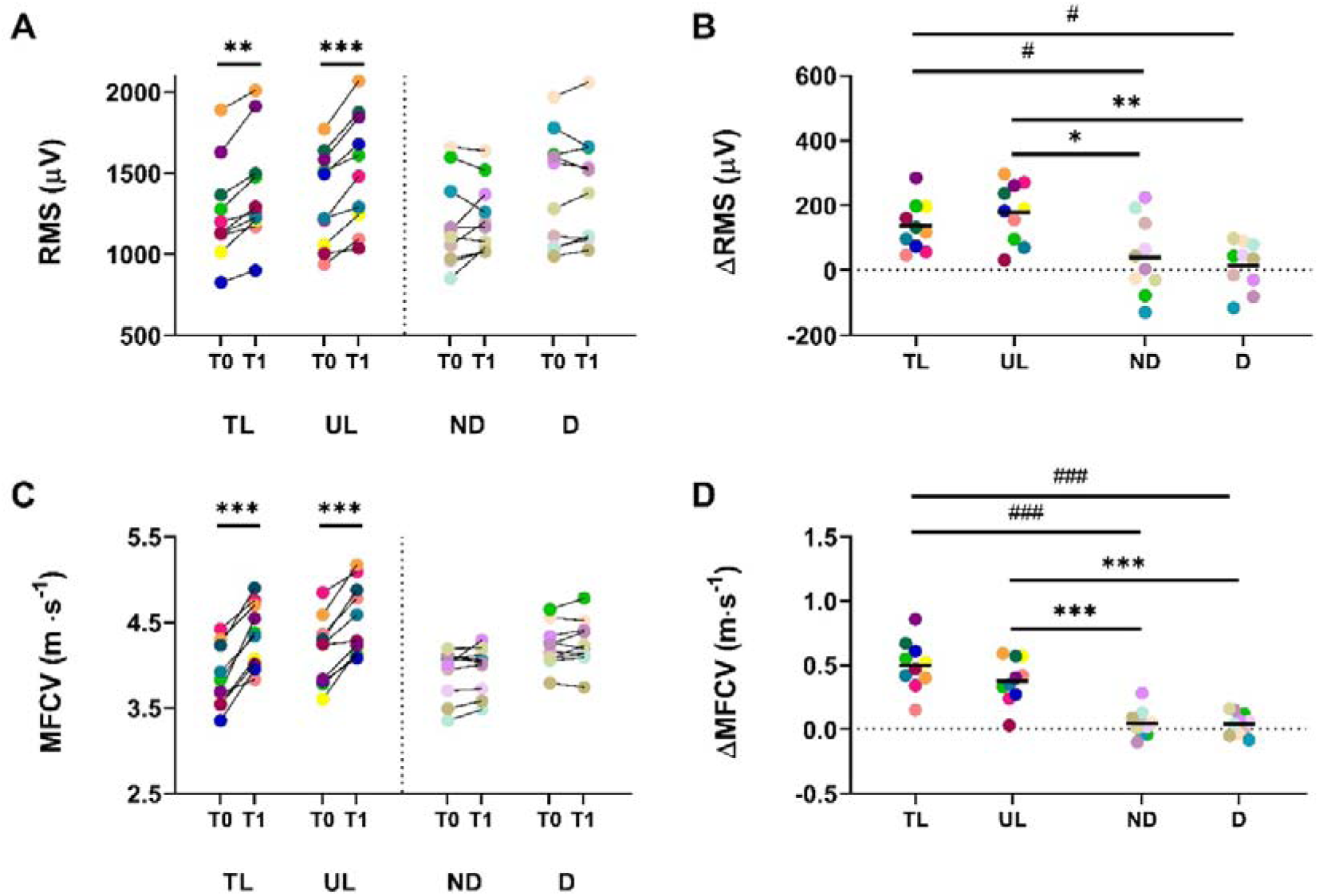
RMS and MFCV adaptations. (A) Absolute EMG amplitude (RMS) and (C) the speed of action potential propagation along muscle fibers (MFCV) are presented for both the intervention (TL, UL) and control (ND, D) groups before (T0) and after (T1) the 4-week protocols. Delta values in RMS (B) and MFCV (D) for each limb are reported. Each color indicates a single participant in the INT (TL, UL) and CNT (ND, D) groups. #, *p* < 0.05; ###, *p* < 0.001 interaction with TL. *, *p* < 0.05; **, *p* < 0.01; ***, *p* < 0.001 interaction with UL.

MFCV showed a significant *group* × *time* interaction for both TL (*X²* = 49.8, *p* < 0.001 [Fig. 3C]) and UL (*X²* = 22.1, *p* < 0.001 [Fig. 3C]). *Post-hoc* analyses revealed increases in the INT group only (TL: ΔMFCV = 0.44 m · s^-1^ [+12%], *p <* 0.001 [Fig. 3D]; UL: ΔMFCV = 0.32 m · s^-1^ [+8%], *p* < 0.001 [Fig. 3D]), with no changes in the CNT group (*p* > 0.05 [Fig. 3D]). Mixed-effect model showed a significant *limb* effect for delta comparisons (*F_3,28.7_* = 24.2, *p <* 0.001), with greater changes in MFCV in TL compared to ND (ΔMFCV = 0.45 m · s^-1^, *p* < 0.001 [Fig. 3D]) and D limbs (ΔMFCV = 0.46 m · s^-1^, *p* < 0.001 [Fig. 3D]), and in UL compared to ND (ΔMFCV = 0.33 m · s^-1^, *p* < 0.001 [Fig. 3D]) and D limbs (ΔMFCV = 0.33 m · s^-1^, *p* < 0.001 [Fig. 3D]).

### MU properties

A significant *group* × *time* interaction was observed for MUDR_MEAN_ in both TL (*X²* = 17.3, *p* < 0.001) and UL (*X²* = 12.3, *p* = 0.002). *Post-hoc* analyses showed increases only in the INT group, with higher MUDR_MEAN_ in TL (ΔMUDR_MEAN_ = 4.6 pps, *p* < 0.001 [Fig. 4A-B]) and UL (ΔMUDR_MEAN_ = 3.5 pps, *p* < 0.001 [Fig. 4A-B]), while no changes were observed in the CNT group (*p* > 0.05 [Fig. 4A-B]). The mixed-effect model for delta values revealed a significant *limb* effect (*F_3,18.6_* = 39.8, *p <* 0.001). Pairwise comparisons revealed larger increases in TL than in UL (ΔMUDR_MEAN_ = 0.72 pps, *p* = 0.02 [Fig. 4C]), ND (ΔMUDR_MEAN_ = 4.43 pps, *p* < 0.001 [Fig. 4C]), and D (ΔMUDR_MEAN_ = 4.24 pps, *p* < 0.001 [Fig. 4C]). UL changes were also greater than those observed in ND (ΔMUDR_MEAN_ = 3.71 pps, *p* < 0.001 [Fig. 4C]) and D limbs (ΔMUDR_MEAN_ = 3.53 pps, *p* < 0.001 [Fig. 4C]).

**Figure 4.**
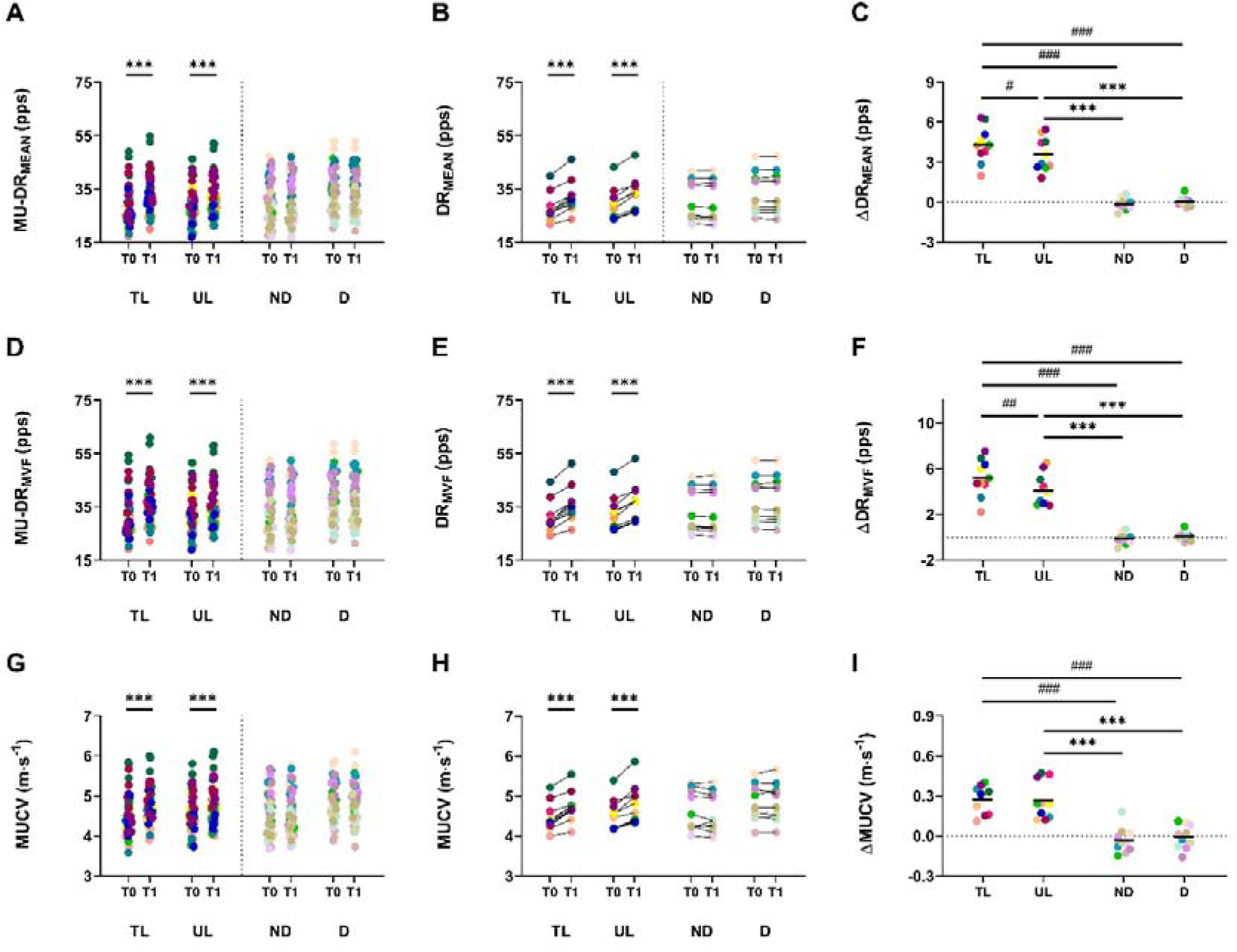
Adaptations in MU discharge characteristics. Mean MU discharge rate results are presented as individual MU data (A), participant average data (B), and *pre-post* delta values (C). MU discharge rate at MVF was also presented as individual MU-data (D), participant-average data (E), and *pre-post* delta values (F). Motor unit conduction velocity is displayed as individual MU-data (G), participant average data (H), and *pre-post* delta values (I). Data for the intervention (TL, UL) and control (ND, D) groups are presented. Each color indicates a single participant in the INT (TL, UL) and CNT (ND, D) groups. ##, *p* < 0.01; ###, *p* < 0.001 interaction with TL. ***, *p* < 0.001 interaction with UL.

A similar observation was found for MUDR_MVF_, with a significant *group* × *time* interaction in TL (*X²* = 14.6, *p* < 0.001) and UL (*X²* = 9.2, *p* = 0.002). *Post-hoc* tests indicated increases only in the INT group (TL: ΔMUDR_MVF_ = 4.9 pps, *p* < 0.001 [Fig. 4C-D]; UL: ΔMUDR_MVF_ = 3.9 pps, *p* < 0.001 [Fig. 4C-D]), whereas CNT showed no changes (*p* > 0.05 [Fig. 4C-D]). The mixed-effect model indicates a significant *limb* effect for delta (*F_3,18.9_* = 44.5, *p <* 0.001). MUDR_MVF_ changes were greater in TL than UL (ΔMUDR_MEAN_ = 1.10 pps, *p* = 0.003 [Fig. 4F]), ND (ΔMUDR_MVF_ = 5.30 pps, *p* < 0.001 [Fig. 4F]), and D (ΔMUDR_MVF_ = 5.10 pps, *p* < 0.001 [Fig. 4F]). Similarly, UL changes exceeded those in ND (ΔMUDR_MVF_ = 4.20 pps, *p* < 0.001 [Fig. 4F]) and D limbs (ΔMUDR_MVF_ = 4.00 pps, *p* < 0.001 [Fig. 4F]).

A significant *group* × *time* interaction was observed for MUCV in both TL (*X²* = 12.2, *p* < 0.001) and UL (*X²* = 12.3, *p* = 0.002). *Post-hoc* analyses showed increases only in the INT group, with higher MUCV in TL (ΔMUCV = 0.3 m · s^-1^, *p* < 0.001 [Fig. 4G-H]) and UL (ΔMUCV = 0.3 m · s^-1^, *p* < 0.001 [Fig. 4G-H]), while no changes were observed in the CNT group (*p* > 0.05 [Fig. 4G-H]). The mixed-effect model for delta values revealed a significant *limb* effect (*F_3,29.5_* = 23.1, *p <* 0.001). Pairwise comparisons revealed larger deltas in TL than in ND (ΔMUCV = 0.31 m · s^-1^, *p* < 0.001 [Fig. 4I]) and D (ΔMUCV = 0.28 m · s^-1^, *p* < 0.001 [Fig. 4I]). UL changes were also greater than those observed in ND (ΔMUCV = 0.30 m · s^-1^, *p* < 0.001 [Fig. 4I]) and D limbs (ΔMUCV = 0.27 m · s^-1^, *p* < 0.001 [Fig. 4I]).

### Associations between neural and force adaptations

Linear regression analyses revealed strong associations between ΔMVF and all neuromuscular parameters in both limbs. Specifically, ΔMVF was strongly associated with ΔRMS (TL: R^2^ = 0.75, *p* = 0.001; UL: R^2^ = 0.75, *p* = 0.001 [Fig. 5A-E]), ΔMFCV (TL: R^2^ = 0.74, *p* = 0.001; UL: R^2^ = 0.77, *p* < 0.001 [Fig. 5B-F]), ΔDR_MEAN_ (TL: R^2^ = 0.71, *p* = 0.002; UL: R^2^ = 0.75, *p* = 0.001 [Fig. 5C-G]), and ΔDR_MVF_ (TL: R^2^ = 0.70, *p* = 0.002; UL: R^2^ = 0.70, *p* = 0.001 [Fig. 5D-H]).

**Figure 5.**
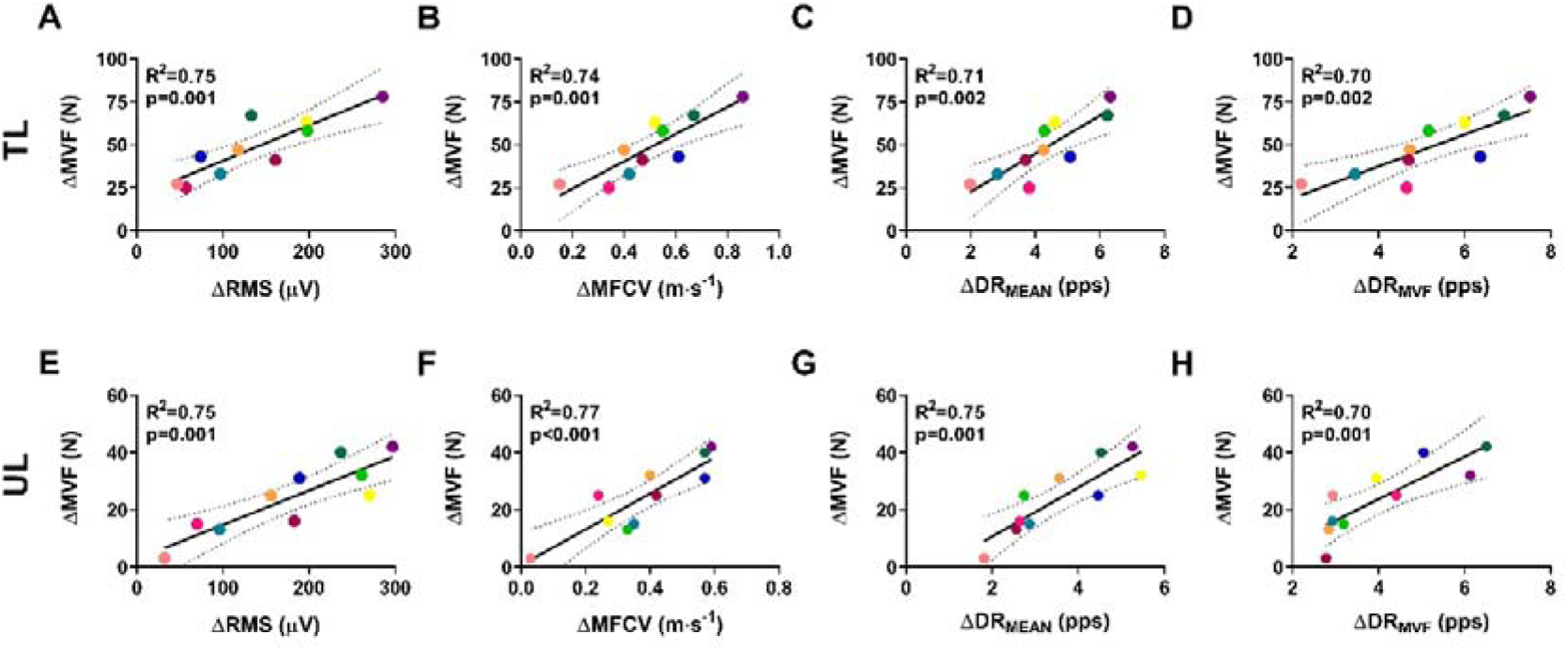
Associations between changes in MVF and neuromuscular parameters. For trained limbs (A-D) and untrained limbs (E-H), the associations between the change in MVF as a function of the change in the neuromuscular adaptations are presented in scatter plots. Each marker refers to a participant’s value. TL, trained limbs; UL, untrained limbs.

Additionally, adaptations in TL and UL were strongly correlated across all assessed variables: ΔRMS (R^2^ = 0.77, *p* < 0.001 [Fig. 6A]), ΔMFCV (R^2^ = 0.77, *p* < 0.001 [Fig. 6B]), ΔDR_MEAN_ (R^2^ = 0.71, *p* = 0.002 [Fig. 6C]), and ΔDR_MVF_ (R^2^ = 0.70, *p* = 0.03 [Fig. 6D]). We also found a significantly strong association between ΔDR_MVF_ and MUCV at baseline in both the TL (R^2^ = 0.50, *p* < 0.001 [Fig. 7A]) and UL (R^2^ = 0.57, *p* < 0.001 [Fig. 7B]), indicating that the greater gains in DR_MVF_ occurred in MU presenting a higher CV. We also observed a significantly strong association between the change in MUCV as a function of the change in discharge rate (ΔMUCV-ΔDR_MVF_) in both the TL (R^2^ = 0.51, *p* < 0.001 [Fig. 7C]) and UL (R^2^ = 0.60, *p* < 0.001 [Fig. 7D]), suggesting that greater changes in DR_MVF_ were accompanied by corresponding greater changes in MUCV for both limbs.

**Figure 6.**
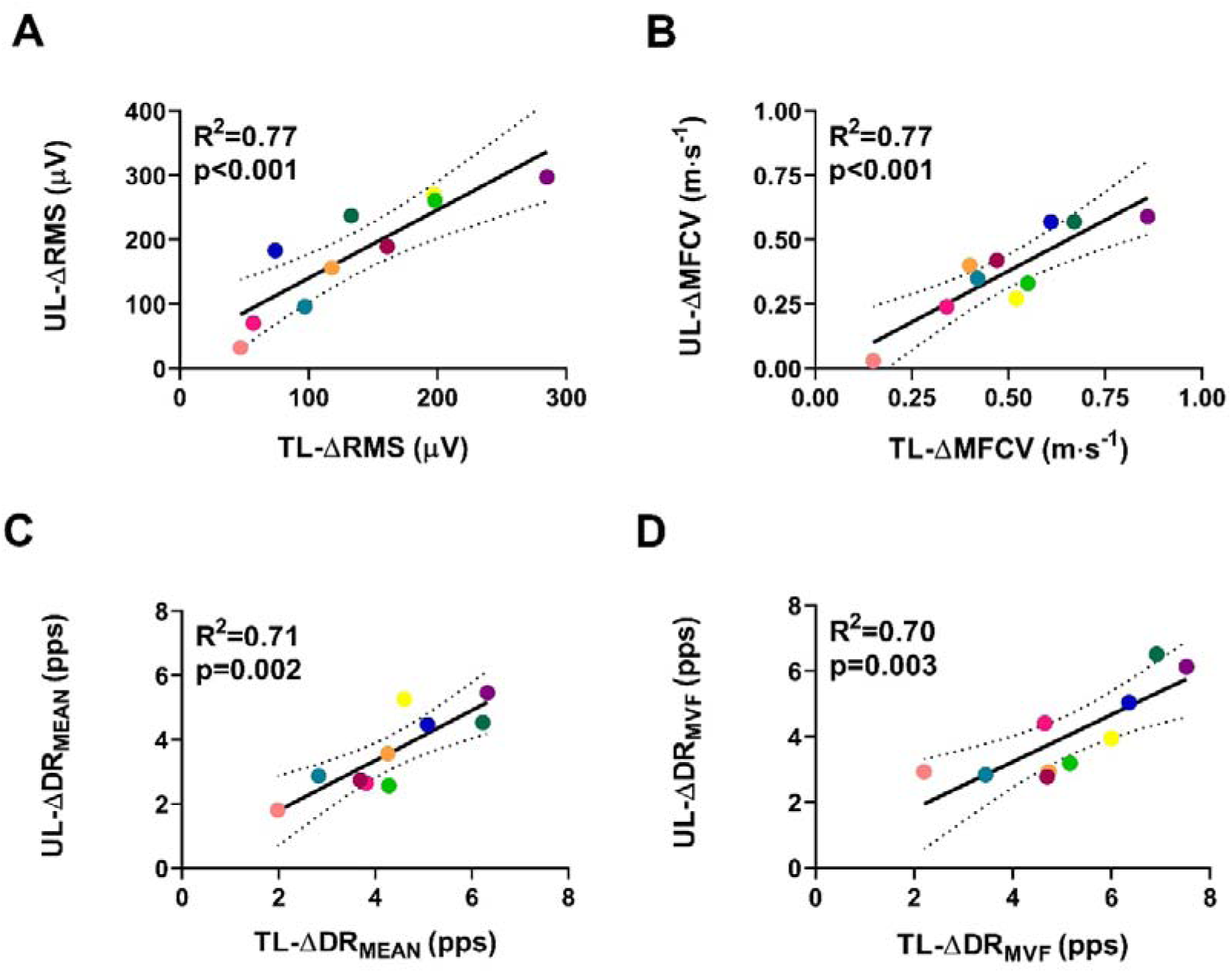
Associations between TL and UL adaptations. Scatter plots show the association between changes in RMS (A), MFCV (B), DR_MEAN_ (C), and DR_MVF_ (D) between TL and UL. The strong association for all neuromuscular variables suggests the transfer of adaptations from the exercised limb to the contralateral untrained side. TL, trained limb; UL, untrained limb.

**Figure 7.**
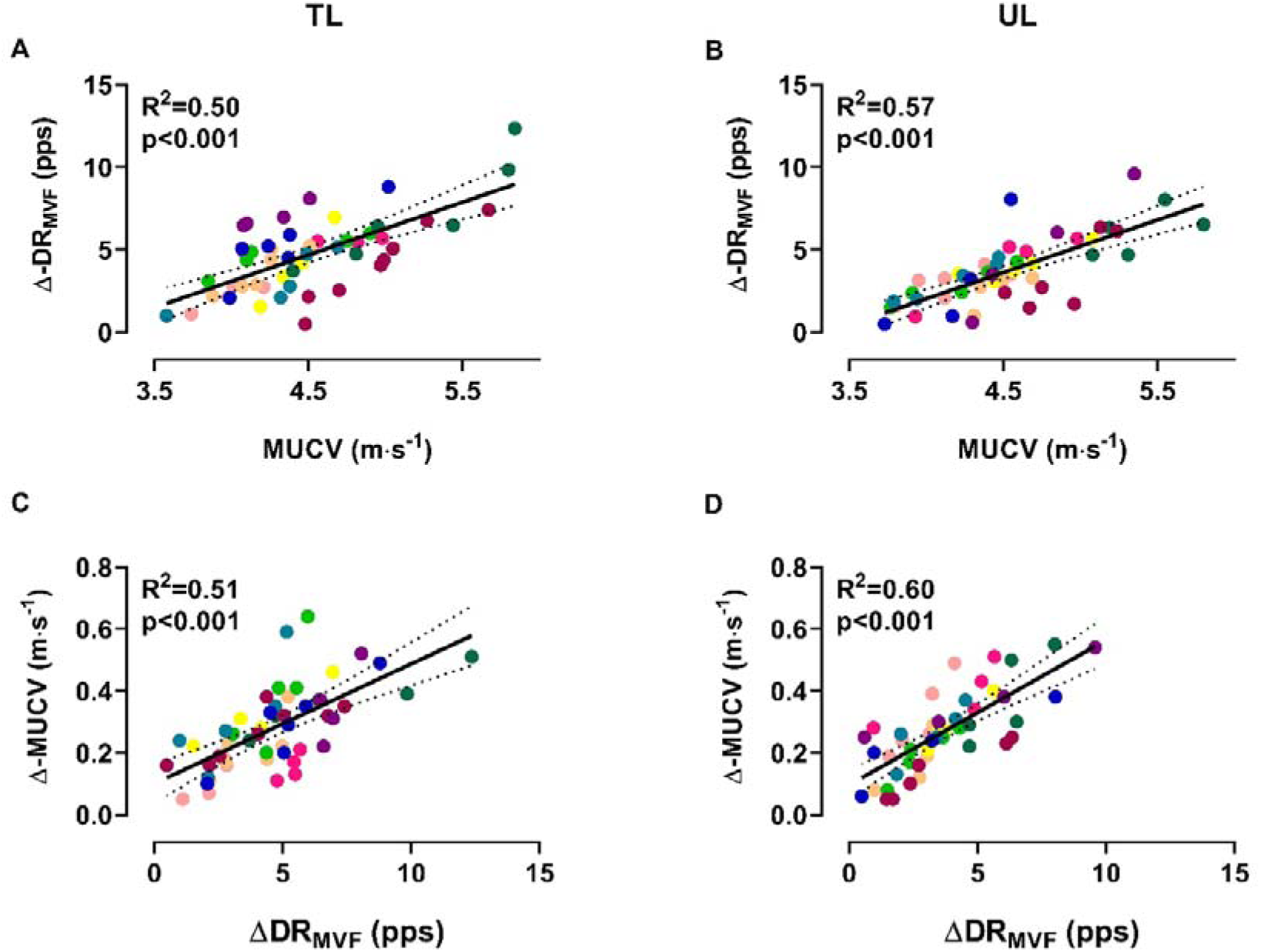
Associations between MUCV and ΔDR_MVF_. Scatter plots show the association between changes in DR_MVF_ as a function of MUCV at *baseline* in both the trained (A) and untrained limb (B). Scatter plots also show the association between *changes* in MUCV and DR_MVF_ for both the trained (C) and untrained (D) limbs. Each marker represents a longitudinally tracked motor unit, and each color refers to a participant. TL, trained limb; UL, untrained limb.

## Discussion

The principal finding of this study is that resistance training induces a differential pattern of MU adaptation during MVC contractions, in which higher-threshold units contribute proportionally more to the increase in neural drive than lower-threshold units. Specifically, the increase in MU discharge rate after training was greater in MUs with higher baseline conduction velocity, indicating that adaptations during maximal force production are not uniformly distributed across the MU pool. During MVCs, further recruitment is limited, and low-threshold units are less able to substantially increase their force contribution because their discharge rates are closer to saturation. Under these conditions, increases in maximal force are expected to depend primarily on enhanced output of higher-threshold spinal motoneurons. Our findings provide direct evidence for this mechanism. A second important result is that these neural adaptations were expressed bilaterally, despite unilateral training. In both the trained and contralateral untrained limbs, maximal contractions after training were characterized by greater surface EMG amplitude, increased MFCV, increased MUCV, and higher MU discharge rates, all consistent with an increased neural drive to the motoneuron pool. The close correspondence between limbs indicates that the neural adjustments supporting gains in maximal force are not confined to the trained side, but are shared bilaterally and likely contribute to cross-education.

These physiological findings were made possible by overcoming a longstanding methodological limitation in the study of resistance training. While MU adaptations have been characterized extensively during submaximal contractions, it has remained difficult to assess the same MUs longitudinally during MVCs because of signal complexity, amplitude cancellation, overlap of action potentials, and the limited duration of maximal-force trials. By longitudinally tracking the same MUs across sessions during MVCs, the present study was able, for the first time, to directly characterize how MU behaviour changes before and after training under maximal force conditions.

This distinction from previous work is important. Earlier studies of cross-education during submaximal contractions indicate that changes in MU recruitment are the main neural mechanism underlying strength transfer to the untrained limb. The present results show that a different mechanism predominates during MVCs, that is enhanced rate coding, particularly in higher-threshold MUs. Thus, the neural adaptations to resistance training depend on the force level at which they are examined, and a full account of training-induced increases in maximal force requires direct investigation of MU behaviour during maximal efforts.

### Reliability of MU decoding from HDsEMG at MVC

To our knowledge, this is the first study to employ MU decomposition and longitudinal tracking across an intervention using maximal voluntary contraction trials. This approach enabled the identification of the same MUs at baseline and following resistance training, overcoming the limitation of sampling different motor units across time points. Accordingly, it provides novel insights into the neural mechanisms underpinning cross-education at the MU level. Approximately 30% of the identified MUs were successfully tracked, consistent with previous studies employing HDsEMG and longitudinal tracking across intervention time points (Martinez-Valdes *et al*., 2017). The high-to-excellent test–retest reliability observed across groups and limbs further supports the accuracy of the tracking procedures, reinforcing the robustness of the present findings (Goodlich *et al*., 2023; Ten Hove *et al*., 2024).

Additionally, because HDsEMG samples a broad muscle region and enables direct MU characterization, RMS and MFCV have been shown to be highly reliable across sessions (Del Vecchio *et al*., 2017; Casolo *et al*., 2020b) and are unlikely to represent sampling artifacts or morphological changes. Furthermore, our findings support previous proposals that increases in EMG amplitude and conduction velocity reported at submaximal contractions extend to maximal efforts in the trained limb (Del Vecchio *et al*., 2025). However, the untrained limb exhibited a markedly different pattern. Whereas previous studies have consistently reported no changes in EMG amplitude during submaximal contractions following unilateral training (Green & Gabriel, 2018), the present study revealed a significant increase in RMS during MVC. This dissociation between submaximal and maximal contraction intensities represents a novel finding and suggests that the neural mechanisms underlying cross-education are intensity-dependent. Specifically, adaptations that are not evident during submaximal force production may become apparent only when the nervous system is required to generate maximal neural drive. Taken together, these findings provide the first evidence that the mechanisms underlying cross-education at maximal contraction differ from those operating at submaximal intensities, highlighting the importance of contraction intensity when interpreting contralateral neuromuscular adaptations. Importantly, this interpretation should be confirmed in future studies, including appropriate control conditions.

### RMS, MFCV, and MUCV adaptations

RMS and MFCV provide global information on the biophysical properties of the active MUs and their cumulative electrical activity during contractions. RMS quantifies the magnitude of the surface EMG signal and, in matched contraction epochs, increases in RMS typically reflect an increased overall MU spiking activity (Del Vecchio *et al*., 2017). Importantly, a higher RMS, together with a higher MUDR, is consistent with an enhanced neural drive to the muscle, even in the absence of peripheral adaptations. This interpretation is supported by recent decomposed EMG studies showing strong within-participant associations between discharge rate and absolute RMS changes (Del Vecchio *et al*., 2025).

Considering the short duration of the training, these early gains in MVF are unlikely to be associated with any increase in muscle thickness, fascicle angle or length (Blazevich *et al*., 2007), cross-sectional area or twitch torque, which are known to occur only after 4 to 5 weeks of regular strength training (Van Cutsem *et al*., 1998; Blazevich *et al*., 2007; Nuzzo *et al*., 2017). In the trained limb, increased MUCV and MFCV are consistent with prior evidence (Cadore *et al*., 2014; Casolo *et al*., 2020a), although the mechanisms underlying these adaptations for short-term RT are subject to ongoing debate. CV is biophysically tied to fiber membrane excitability and fiber diameter, and is sensitive to factors such as the resting membrane potential and the activity or density of Na⁺/K⁺-ATPase pumps (Clausen, 2003; Christiansen, 2019). However, changes in CV are not purely peripheral, because single-MU CV is also influenced by discharge rate (the so-called *velocity-recovery* function of muscle fibers) (Stalberg, 1966; Campanini *et al*., 2009). The increase in MFCV observed here cannot reflect a different MU population, since the full pool was presumably recruited at maximal force across all conditions, and the training duration was not sufficient to affect membrane properties (Lecce *et al*., 2026). Therefore, it is likely that the changes in MFCV reflected an increase in MUCV, driven by higher DR. The relative change in CV with DR for the biceps brachii muscle has previously been reported to be approximately 1.3%/pps (Nishizono *et al*., 1989), which means that CV increases by 1.3% for each increase in DR by 1 pps. These previous results are well in agreement with the changes in MUCV and DR documented in this study. Figure 4 indeed shows that an average change of approximately 5 pps corresponded to an increase of 0.3 m·s^-1^ in MUCV. Assuming a mean CV of ∼ 4 m·s^-1^ (see *supplementary statistical table*), an increase by 0.3 m·s^-1^ corresponds to a 7.5% increase, which, over a range of 5 pps, leads to 1.5%/pps. This explains the changes in MFCV and MUCV with the *velocity-recovery* function of muscle fibres without any training-induced change in membrane fibre properties.

In addition, at the level of individual motor units, there was a significant linear association between the training-induced increase in MUDR and the baseline MUCV (i.e., the faster the baseline CV for a given MU) the larger the increase in its DR following training. Because CV is directly associated with fiber size (Casolo *et al*., 2023) and, therefore, with the recruitment threshold (Andreassen & Arendt-Nielsen, 1987), the higher-threshold MUs experienced the largest increase in DR following training (Fig. 7). This indicates a greater contribution of higher-threshold MUs to the increase in neural drive and, ultimately, to the greater MVF. Interestingly, an increase in the contribution of high-threshold motor units has also been reported as the main mechanism of training-induced MVF for submaximal contractions (Del Vecchio *et al*., 2019a; Casolo *et al*., 2020a). However, while this is achieved by compression of recruitment thresholds in submaximal efforts, during maximal efforts, an increase in high-threshold MU activity is accompanied by a differential increase in DR between MUs with different thresholds, likely due to partial DR saturation in low-threshold units. Notably, these adaptations were also present in the untrained muscles, where hypertrophic changes are even less likely to occur (Phillips, 2000; Mirto *et al*., 2025).

Taken together, the concurrent increase in RMS, MFCV, and MUCV during maximal contractions suggests a similar pool of concurrently active MUs, but with a different distribution of discharge rates and a relatively greater contribution from higher-threshold units whose action potentials exert a stronger influence on the average surface estimate.

### Neural drive adaptations

The discharge rate across a relatively large sample of MUs provides an estimate of neural drive to muscle (Farina *et al*., 2014; Negro *et al*., 2016b; Del Vecchio *et al*., 2020; Lecce *et al*., 2025b). In the present study, we observed an increase in DR_MVF_ of ∼5 pps (∼4 pps as DR_MEAN_) in both the trained and untrained limbs. In the trained limb, this increase appears greater than that previously reported after resistance training in the tibialis anterior muscle (∼3 pps) using an isometric ballistic plus sustained contraction-based training program (Del Vecchio *et al*., 2019a), as well as in our previous investigation in the biceps brachii muscle, but during submaximal contractions, where increases were closer to ∼2.5 pps (Lecce *et al*., 2025a). These comparisons, however, are only possible for the trained limb, as previous studies reported no changes in discharge rate in the untrained limb during submaximal contractions (Green & Gabriel, 2018; Lecce *et al*., 2025c, 2025a). The larger increase observed here may therefore reflect both differences in the training stimulus and the contraction intensity used to assess neural adaptations. Indeed, whereas previous studies quantified discharge rate during submaximal contractions (up to 70% MVF), the present study assessed *maximal voluntary contractions*, likely providing greater sensitivity to detect adaptations in high-threshold MUs and supraspinal drive mechanisms that remain concealed at submaximal contraction intensities.

Higher DR_MEAN_ indicates that, on average, the MUs discharge at a higher frequency throughout the contraction, supporting greater force output (Del Vecchio *et al*., 2019a). Besides, higher DR_MVF_ indicates a greater peak discharge rate at maximal force and, therefore, a greater capacity to generate maximal muscle force (Lecce *et al*., 2025b). Higher discharge rates promote twitch fusion and increase contractile output (Fuglevand *et al*., 1993; Macefield *et al*., 1996; Heckman & Enoka, 2012; Caillet *et al*., 2022). Accordingly, the observed gains in maximal strength are plausibly explained by MU adaptations, including increased discharge rates of higher-threshold MUs (Del Vecchio *et al*., 2019a; Škarabot *et al*., 2021; Lecce *et al*., 2026), as supported by the strong associations between changes in these variables and MVF (Fig. 5).

These modifications were observed on both the trained and untrained sides, contrasting with prior studies conducted during submaximal contractions that reported no changes in discharge rate following unilateral training (Lecce *et al*., 2025c, 2025a). During submaximal contractions, the central nervous system preferentially recruits low- to mid-threshold MUs and may insufficiently engage the high-threshold population or the neural processes that become increasingly important during maximal efforts (Fuglevand *et al*., 1993; Deluca & Erim, 1994). By examining maximal voluntary contractions, we therefore identified rate-coding adaptations that are not apparent at lower contraction intensities. These changes may reflect adaptations occurring at multiple levels of the neuromuscular system, including increased corticospinal excitability, reduced intracortical or interhemispheric inhibition, enhanced synaptic input to motoneurons, reduced presynaptic inhibition of Ia afferents, or alterations in motoneuronal intrinsic properties (Škarabot *et al*., 2021; Lecce *et al*., 2026). The bilateral increase in MUDR is consistent with neural adaptations that are not restricted to the trained limb and may involve both supraspinal and spinal mechanisms (Ruddy & Carson, 2013; Hendy & Lamon, 2017; Carson, 2020). However, because the present study did not directly assess the relative contribution of these pathways, the specific mechanisms responsible for the observed changes cannot be determined. These adaptations are particularly relevant at near-maximal effort, because once the pool of available motor units is fully recruited (∼ 90% MVF in the biceps brachii) (Kukulka & Clamann, 1981), additional gains in muscle force can only be achieved by increasing the discharge rate (Heckman & Enoka, 2012).

Resistance training (both bilateral and unilateral) increases net excitatory drive to the motoneuron pool (Glover & Baker, 2020; Škarabot *et al*., 2021; Lecce *et al*., 2026). In this context, lower-threshold motor units already operate near their discharge ceiling, whereas higher-threshold units retain a larger dynamic range for rate modulation and can therefore increase their firing rates more under the same enhanced drive (Heckman & Enoka, 2012; Enoka & Duchateau, 2017). This interpretation is supported by the significant association between the change in DR_MVF_ and baseline MUCV (Fig. 7), indicating that the greatest increases in discharge rate during maximal voluntary force occur in higher-threshold MUs.

### Cross-education of maximal muscle force

The parallel increases in RMS, MFCV, and MUDR (Fig. 6) in both limbs indicate that unilateral resistance training enhanced maximal force primarily through greater neural drive to the motoneuron pool. Because motor unit recruitment in biceps brachii is expected to be largely complete near maximal force, these adaptations are more consistent with altered rate-coding behavior within the active motor unit pool than with additional recruitment. The association between baseline MUCV and the increase in discharge rate further suggests that higher-threshold motor units (Casolo *et al*., 2023) contributed more to the increase in force output. This is physiologically consistent with the fact that lower-threshold MUs are already closer to their maximal discharge rate during maximal efforts, whereas higher-threshold MUs retain greater capacity for further increases in discharge rate (Macefield *et al*., 2000; Enoka & Duchateau, 2017).

Importantly, the untrained limb exhibited similar neural adaptations despite the absence of direct mechanical loading (Lecce *et al*., 2026), supporting the neural origin of cross-education. Unlike submaximal contractions, where strength transfer appears mainly related to altered recruitment (Lecce *et al*., 2025c, 2025a), maximal-force transfer was characterized by enhanced discharge rates, indicating that cross-education at high contraction intensities depends predominantly on increased descending excitatory input to spinal motoneurons. This may reflect a greater corticospinal contribution and reduced intracortical inhibition as voluntary effort rises (Hammond & Vallence, 2007; Castelli *et al*., 2025), potentially supplemented by greater brainstem drive that further increases the net descending excitatory input with rising effort (Glover & Baker, 2022), thereby increasing the effective neural drive to muscle as contraction intensity increases (Škarabot *et al*., 2022, 2025).

These findings support the view that unilateral resistance training induces bilateral adaptations within shared motor networks (Ruddy & Carson, 2013; Ruddy *et al*., 2017). increasing the capacity of both limbs to generate maximal force through enhanced motoneuron output, particularly from higher-threshold motor units (Farthing *et al*., 2007; Ruddy & Carson, 2013; Hendy & Lamon, 2017; Calvert & Carson, 2022; Lecce *et al*., 2026).

## Limitations and future directions

Although we observed parallel changes in MFCV, EMG amplitude, and estimates of neural drive, several limitations should be considered. First, the maximality of the MVCs was not verified using techniques such as twitch interpolation; therefore, changes in neural drive during maximal efforts should be interpreted cautiously. Second, antagonist muscle activity was not recorded. Consequently, increases in elbow flexion force may have been influenced by changes in triceps coactivation in addition to changes in agonist activation. Third, although the observed adaptations are consistent with neural mechanisms, we did not directly assess corticospinal, intracortical, interhemispheric, or spinal function. Therefore, the relative contribution of supraspinal and spinal pathways cannot be determined. Measures of EMG amplitude alone also do not allow the respective contributions of MU recruitment and discharge rate to be distinguished without decomposition of the interference EMG signal.

Furthermore, both MFCV and EMG amplitude can only be interpreted as indicators of neural adaptations in the absence of substantial muscular changes. Although structural adaptations are unlikely after a short unilateral intervention and are not typically expected in the untrained limb, changes in muscle morphology or volume-conductor properties could have influenced these measures. Similarly, while MFCV is commonly used as an indirect indicator of MU recruitment, it should not be considered a direct measure of recruitment threshold. Finally, the number of longitudinally tracked motor units was relatively small and may not fully represent adaptations across the entire motor-unit pool. Future studies combining assessments of MU behavior, antagonist coactivation, voluntary activation, central and spinal neural function, and muscle morphology are needed to further clarify the mechanisms underlying cross-education of maximal muscle force.

## Conclusions

Unilateral resistance training increased MU discharge rate, RMS, and MFCV during maximal voluntary contractions in both the trained and contralateral untrained limbs. The increase in discharge rate was greater in MUs with higher baseline conduction velocity, indicating a greater contribution of higher-threshold motor units to the increased neural drive to muscle. This greater rate coding adaptation for higher-threshold units likely reflects their lower initial firing-rate saturation compared with lower-threshold units, which operate closer to their firing-rate ceiling at maximal effort. In the trained limb, adaptations were similar across maximal and submaximal contractions. By contrast, the untrained limb showed an intensity-dependent pattern: submaximal force increases were mainly associated with increased MU recruitment, whereas maximal force increases were primarily driven by higher discharge rates of active MUs. Collectively, these findings show that cross-education of maximal strength is mediated by recruitment and rate-coding adaptations, with maximal-force gains in the untrained limb driven primarily by a preferential increase in the firing of higher-threshold MUs.

## Additional information

### Author contributions

E.L. and I.B. conceived and designed the study. E.L., P.A. and I.B. collected data. Data analysis and interpretation were collaboratively carried out by E.L., P.A., A.D.V., A.C., F.F., D.F. and I.B. All authors contributed to manuscript drafting, critical revisions, edits and the overall refinement of the text. All authors listed meet the criteria for authorship, have made substantial contributions to the work, and have reviewed and approved the final manuscript. All authors agree to be accountable for all aspects of the work.

## Data availability statement

The data supporting the findings of this study and MATLAB scripts used for data analysis are available upon reasonable request from the corresponding author.

## Competing interests

None declared.

## Funding

The present work was supported by the grant - [*CDR2.DIP2025 – prot. 007693*].

## Notes

### Competing Interest Statement

The authors have declared no competing interest.

